# eIF4G2-dependent translation restrains pancreatic cancer progression

**DOI:** 10.64898/2026.03.14.711799

**Authors:** Justin Powers, Wei Lai, Hiroki Kobayashi, Alvaro Curiel-Garcia, Pardis Ahmadi, Elizabeth Valenzuela, Marko Jovanovic, Alejandro Chavez, Iok In Christine Chio

## Abstract

Pancreatic ductal adenocarcinoma (PDA) is among the most lethal cancers, driven by cellular plasticity that fuels therapeutic resistance and early dissemination. The contribution of translational control to this plasticity remains poorly understood. Through an *in vivo* CRISPR/Cas9 screen, we identify the non-canonical initiation factor eIF4G2 (DAP5/NAT1) as a translational checkpoint restraining PDA progression. Loss of eIF4G2 accelerated tumor growth, induced poorly differentiated, basal-like histology, and triggered widespread metastasis. Ribosome profiling revealed that eIF4G2 loss does not alter bulk protein synthesis but instead impairs translation of a selective regulon, including tumor suppressors such as PTEN and CREBBP. Functional studies confirmed that PTEN loss was sufficient to drive dedifferentiation but insufficient to promote metastasis, implicating the broader eIF4G2-dependent program, including translational control of transcriptional regulators like CREBBP, in limiting dissemination. Consistently, eIF4G2-deficient tumors exhibited transcriptomic enrichment of programs related to migration and wound healing. Computational inference from human PDA datasets revealed reduced eIF4G2 activity in metastases, aligning with basal-like features and predicting poorer survival. These results support a model in which eIF4G2 maintains epithelial identity and restrains metastatic potential, highlighting selective translation as a determinant of PDA subtype and clinical outcome.

## INTRODUCTION

Pancreatic ductal adenocarcinoma (PDA) remains one of the most lethal human malignancies, with a five-year survival rate below 12% despite advances in surgery, systemic therapy, and supportive care (1). Poor outcomes are driven by late diagnosis, intrinsic therapeutic resistance, and early metastatic spread (2,3). Genetically, PDA is relatively homogeneous: activating mutations in *KRAS* occur in the vast majority of tumors, while *TP53, CDKN2A*, and *SMAD4* are inactivated in large subsets (4,5). These alterations provide a common oncogenic backbone upon which extensive phenotypic heterogeneity emerges. Over the past decade, transcriptomic studies have organized this heterogeneity into clinically meaningful subtypes. Tumors segregate along an epithelial “Classical” axis, marked by well-differentiated glandular morphology, epithelial gene programs, and relatively better outcomes, versus a “Basal-like” (or squamous) axis characterized by dedifferentiation, mesenchymal features, therapeutic refractoriness, and poor prognosis (6). This framework has clarified important differences in disease behavior and treatment response (7), yet the molecular mechanisms that permit plasticity between these states remain incompletely understood. In particular, while transcriptional reprogramming has been extensively studied, the role of translational checkpoints in governing PDA cell states has remained unexplored *in vivo*.

Protein synthesis is among the most energy-consuming cellular processes, accounting for ~20% of energy usage in normal cells and up to 40% in cancer (8). Accordingly, oncogenic signaling pathways, including PI3K/AKT/mTOR and MEK/MNK, often converge on the translation machinery (9). Many of these pathways impinge on the eIF4F complex, the central driver of cap-dependent translation (10). The eIF4F complex consists of the cap-binding protein eIF4E, the scaffolding protein eIF4G1, and the RNA helicase eIF4A (11). Elevated eIF4F activity promotes the translation of oncogenic transcripts and correlates with poor prognosis and therapy resistance across various cancers (9,12,13). Indeed, we and others have shown that inhibition of eIF4A, the helicase component of the eIF4F complex, strongly impairs PDA cell growth compared to normal ductal cells (14–16). While canonical eIF4F-dependent translation has been intensively studied, far less is known about non-canonical initiation factors. One such factor, eIF4G2 (also known as DAP5 or NAT1), is a paralog of eIF4G1 but lacks the domain that eIF4G1 uses to interact with the cap-binding protein eIF4E (17). Several models have been proposed for the function of eIF4G2, including IRES-mediated (18), m(6)A-mediated (19), and eIF3d-mediated (20) translation initiation. However, these models are based largely on *in vitro* or cultured cell studies, and the role of eIF4G2 *in vivo* has yet to be clarified. Here, we identify eIF4G2 as a translational checkpoint in PDA. Loss of eIF4G2 in PDA cells drives dedifferentiation, “Basal-like” reprogramming, and widespread metastasis through impaired translation of tumor-suppressive regulators, including *Pten* and *Crebbp*. Computational analysis further revealed that reduced eIF4G2 activity characterizes metastatic human PDA and predicts poor patient survival, supporting eIF4G2 as a key barrier to the progression and metastatic dissemination of PDA.

## RESULTS & DISCUSSION

### Genome-wide CRISPR screen identifies eIF4G2 as a suppressor of PDA

To identify genes essential for pancreatic cancer growth *in vivo*, we performed a genome-wide CRISPR knockout screen using a murine KPC (*Kras*^*G12D*^*;Trp53*^*R172H*^*;Pdx1-Cre*) PDA cell line (ccmT2) subcutaneously injected into immunocompetent C57Bl/6J mice (Fig. 1A). The screen was performed by transducing ccmT2 with the Brie mouse whole-genome CRISPR knockout library. Transduced cells were selected using puromycin for 3 days. Following puromycin selection, a baseline sample was collected (Day 0). Cells were then passaged *in vitro* for 17 days to deplete gRNAs targeting common essential genes, at which point an engraftment-ready cell sample was collected (Day 17). Endpoint tumors were harvested 10 days post-injection after reaching an average volume of approximately 1.5 cm^3^. Finally, sgRNA abundance was quantified via deep sequencing from the Day 0, Day 17, and endpoint tumor samples. We confirmed adequate library representation (Supplementary Fig. 1A) and robust depletion of essential genes by Day 17 (Supplementary Fig. 1B), which correlated with DepMap essentiality scores (Fig. 1B). We find that the initial *in vitro* passage successfully depleted common essential genes from the library, resulting in their less pronounced depletion in the tumors (Fig. 1C, Supplementary Fig. 1B) (21). Importantly, our *in vivo* screen successfully identified known tumor suppressors such as *Cdkn2b, Nf2, Pten, Rb1*, and *Smad4*, as well as canonical oncogenes including *Kras* and *Myc* (Fig. 1D). Our results also strongly correlate with an independent screen conducted on pancreatic cancer cells using a metabolic library (Supplementary Fig. 1C) (22), further confirming the technical robustness of our screen. Among the top enriched hits, we identified *Eif4g2*, a non-canonical translation initiation factor. Multiple independent sgRNAs targeting *Eif4g2* were consistently enriched across both the *in vitro* and *in vivo* arms (Fig. 1D and 1E; Supplementary Fig. 1E and 1F), indicating that *Eif4g2* loss provides a selective advantage during tumor growth.

**Figure 1.**
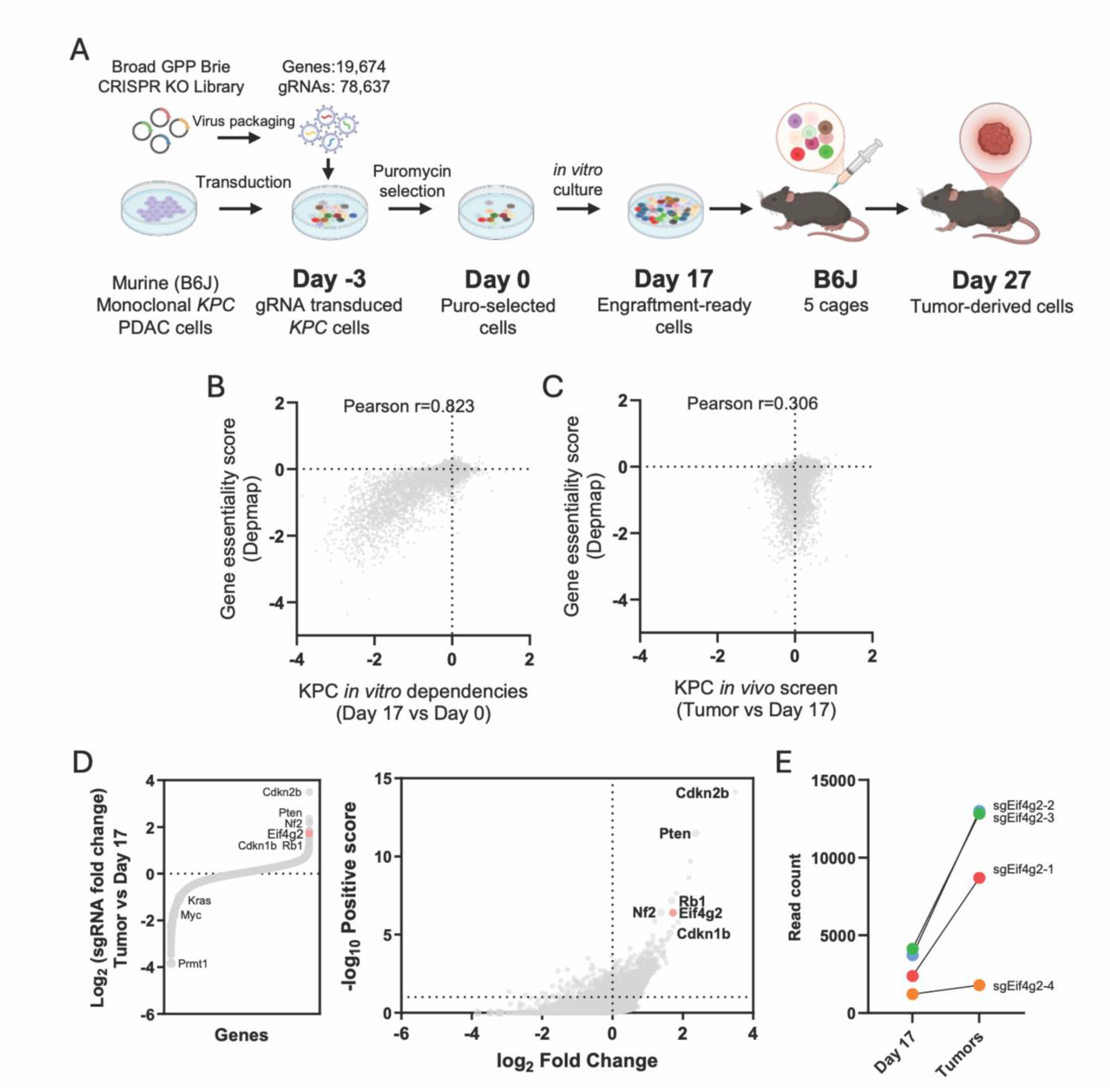
A genome-wide *in vivo* CRISPR screen nominates eIF4G2 as a suppressor of PDA. (A) Schematic of the in vivo CRISPR-Cas9 screening workflow, from library transduction of KPC PDAC cells to tumor harvest in 5 cages of 10-week-old mice. (B) Correlation of gene essentiality scores from the in vitro portion of the screen (beta scores from MAGeCK-MLE, Day 17 vs. Day 0) with pan-cancer essentiality scores from the DepMap database. (C) Reduced correlation of in vivo gene essentiality scores (beta scores, endpoint tumors vs. Day 17) with DepMap scores. (D) Changes in gRNA enrichment between the in vitro outgrowth (Day 17) and in vivo tumors, known tumor suppressors (e.g., *Cdkn2b, Pten*) and oncogenes (e.g., *Kras, Myc*) are highlighted. (E) Normalized read counts for four independent gRNAs targeting *Eif4g2*, showing their strong enrichment in endpoint tumors relative to the Day 17 engraftment-ready cell population. Data are represented as the median of all primary tumors.

eIF4G2 has been linked to selective translation of transcripts involved in stress adaptation and survival (23), but its role in cancer remains unsettled. While some studies suggest oncogenic functions (24,25), others report tumor-suppressive roles (26), and recurrent *EIF4G2* mutations have been observed in tumor genomes (27). Thus, to investigate the role of eIF4G2 in PDA progression, we targeted *Eif4g2* expression by CRISPR-Cas9, using two independent sgRNAs, in murine KPC pancreatic cancer cells (ccmT4, a different KPC line from the ccmT2 screening line). To assess tumorigenic potential *in vivo*, we orthotopically implanted control (sg*Rosa*) or sg*Eif4g2* KPC cells into the pancreas of syngeneic mice. At 5 weeks post-implantation, mice were euthanized for endpoint analyses. Animals harboring *Eif4g2*-deficient tumors (Fig. 2A) exhibited a significant increase in tumor burden, with an average 2-fold increase in tumor volume relative to control (Figs. 2B). sg*Eif4g2* tumors also exhibited elevated mitotic activity, as indicated by increased phospho-Histone H3 (pH3) staining (Fig. 2C). Blinded histopathological analysis of primary tumors revealed a profound shift in differentiation status. While control tumors generated by sg*Rosa* cells ranged from well-to moderately differentiated histological grades (Figs. 2D and 2E), with approximately half displaying glandular structures (Fig. 2F), sg*Eif4g2* tumors were poorly differentiated, exhibiting marked loss of glandular features (Figs. 2D-2F). Strikingly, this dedifferentiated state was accompanied by widespread metastasis (observed in 100% of sg*Eif4g2* tumors; Fig. 2G, Supplementary Table 1). Together, these findings reveal eIF4G2 as a previously unrecognized regulator of PDA, acting to limit tumor growth, maintain epithelial differentiation, and restrain metastatic dissemination. These observations prompted us to investigate whether eIF4G2 enforces these effects through a selective translational program.

**Figure 2.**
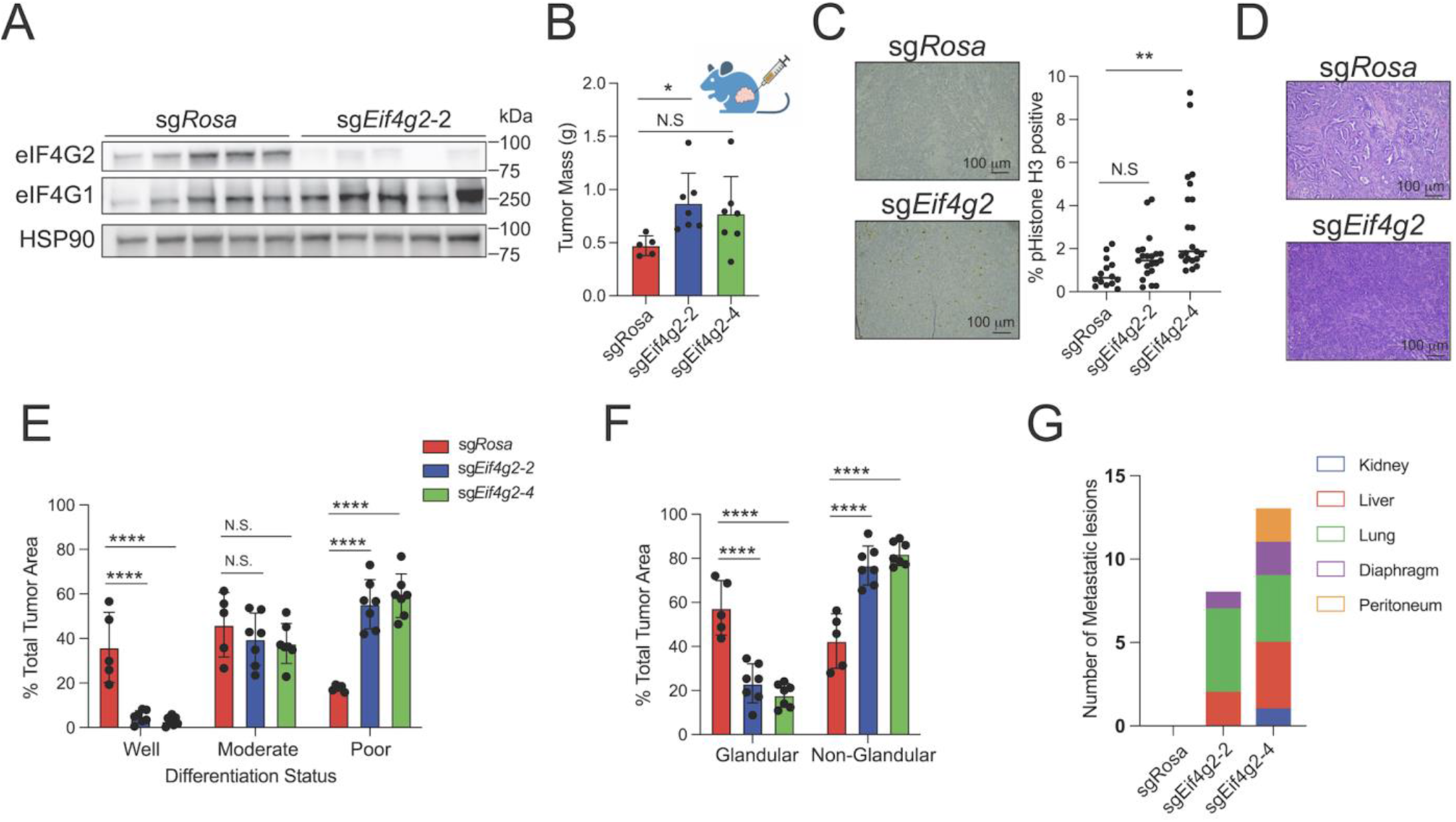
eIF4G2 restrains tumor growth, maintains differentiation, and limits metastasis *in vivo*. (A) Immunoblot analysis of eIF4G2 expression in bulk tumor lysates. (B) Mass of pancreatic tumors at the 5-week endpoint after orthotopic implantation of control (sg*Rosa*) and eIF4G2-deficient (sg*Eif4g2*) KPC cells. sg*Eif4g2-2* and sg*Eif4g2-4* are two sgRNAs targeting *Eif4g2*. (C) Immunohistochemical staining and quantification of phospho-histone H3 in orthotopic PDA tumors. (D-F) Representative H&E images (D) and pathological assessments of sg*Rosa* and sg*Eif4G2* tumor sections based on differentiation status (E) and glandular morphology (F). (G) Number of metastatic lesions across different organ sites. Error bars in this Fig. are means ± SDs. Student’s t-test was performed. ns (not significant) for P ≥ 0.05, * (one asterisk) for P < 0.05, ** (two asterisks) for P < 0.01, *** (three asterisks) for P < 0.001, and **** (four asterisks) for P < 0.0001.

### eIF4G2 drives a selective tumor-suppressive translatome in PDA

In PDA, increased eIF4F activity, particularly through its helicase, eIF4A, promotes tumor growth by enabling the translation of structured, oncogenic mRNAs (9,14-16). Because eIF4G2 lacks the N-terminal eIF4E-binding domain found in its paralog, eIF4G1, but retains the ability to interact with eIF4A, we initially hypothesized that eIF4G2 loss might liberate eIF4A, thereby enhancing eIF4F-dependent, global translation, potentially explaining the increased proliferation of sg*Eif4g2* PDA cells (Supplementary Figs. 2A, 2B). However, we observed no increases in bulk translation, polysome loading, or mTORC1 activity (Supplementary Figs. 2C–2H) upon eIF4G2 loss. These results indicate that eIF4G2 loss likely does not act through canonical eIF4F-dependent programs, prompting us to investigate selective translational changes by ribosome profiling.

To define the selective translational program regulated by eIF4G2, we performed ribosome profiling (Ribo-seq) in three independent murine PDA cell lines (ccmT1, ccmT2, and ccmT4) transduced with sg*Rosa* or sg*Eif4g2*. Libraries met the expected quality metrics (Supplementary Figs. 3A and 3B), with no eIF4G2-dependent changes in elongation dynamics (Supplementary Fig. 3C) and codon usage (Supplementary Fig. 3D). Integration of ribosome-protected fragments (RPF) and mRNA abundances identified 83 transcripts with significantly altered translation efficiency (adjusted p < 0.05), of which 79 were decreased and 4 were increased (Fig. 3A and Supplementary Table 2) in sg*Eif4g2* cells. Among the most strongly downregulated transcripts was *Pten*, a well-established tumor suppressor. Notably, PTEN mutations are rare in human PDA (28), but decreased PTEN protein expression and elevated PI3K-AKT signaling are frequently observed and associated with aggressive disease (29–31), and *Pten* deletion cooperates with mutant *Kras* to accelerate PDA development in experimental models of PDA (32). These findings suggest that translational regulation by eIF4G2 may provide a mechanistic explanation for the loss of PTEN in PDA. Indeed, polysome fractionation confirmed reduced association of *Pten* mRNA within translation-active heavy polysomes (Supplementary Fig. 3E) and decreased PTEN protein levels in both murine (Fig. 3B and Supplementary Fig. 3F) and patient-derived (Fig. 3C) PDA models lacking EIF4G2. To test whether PTEN loss functionally accounts for the phenotype of *Eif4g2* loss, we orthotopically transplanted sg*Pten* KPC cells (Supplementary Fig. 3G). These tumors were significantly larger (Supplementary Fig. 3H) and exhibited poorly differentiated histology (Figs. 3D and 3E), recapitulating the growth and dedifferentiation phenotype of sg*Eif4g2* tumors. However, unlike sg*Eif4g2* tumors, in this same orthotopic model, sg*Pten* tumors did not give rise to metastatic lesions. Thus, while *Pten* translational suppression contributes to enhanced growth and dedifferentiation, additional eIF4G2-dependent targets are likely required to explain the metastatic phenotype. Collectively, these data indicate that eIF4G2 supports a selective, tumor-suppressive translational program that includes PTEN and additional regulators, acting to maintain differentiation and restrain PDA progression.

**Figure 3.**
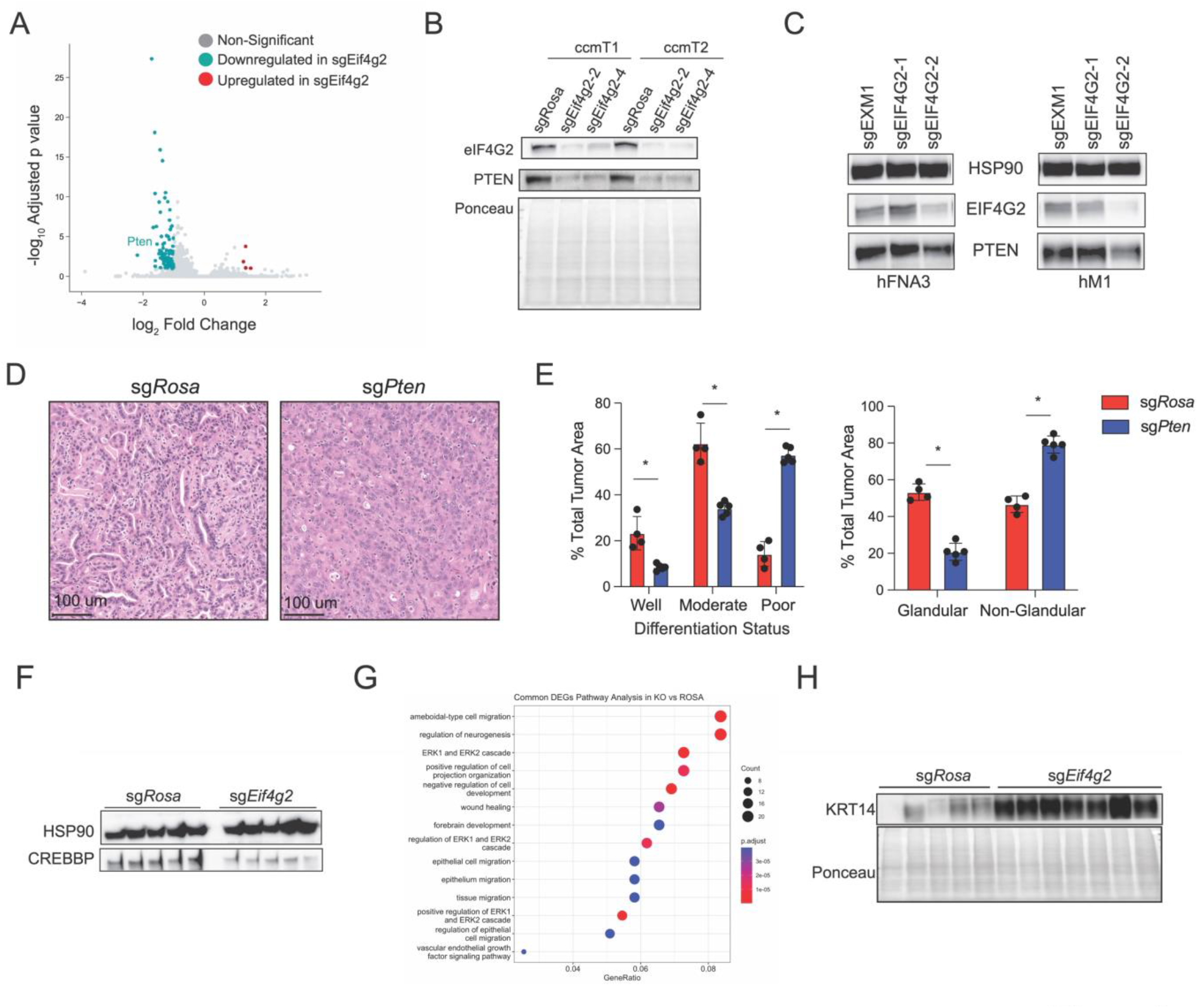
Loss of eIF4G2 impairs a tumor-suppressive translational program. (A) Volcano plot showing changes in translation efficiency in sg*Eif4g2* vs sg*Rosa* KPC cells. (B) Immunoblot analysis of PTEN expression in control (sg*Rosa*) and eIF4G2-deficient (sg*Eif4g2*) KPC cells. (C) Immunoblot analysis of PTEN expression in two independent control (sg*Rosa*) and eIF4G2-deficient (sg*Eif4g2*) patient-derived PDA cell lines. (D) Representative H&E images from sg*Rosa* and sg*Pten* tumor sections. (E) Pathological assessment of sg*Rosa* and sg*Pten* tumor sections based on glandular features and differentiation status. (F) Representative immunoblots of CREBBP in control (sg*Rosa*) and eIF4G2-deficient (sg*Eif4g2*) KPC cells. (G) Gene set enrichment analysis of transcripts upregulated upon eIF4G2 loss. (H) Representative immunoblot analysis of KRT14 expression in control (sg*Rosa*) and eIF4G2-deficient (sg*Eif4g2*) bulk tumor lysates. Error bars in this Fig. are means ± SDs. Student’s t-test was performed. ns (not significant) for P ≥ 0.05, * (one asterisk) for P < 0.05, ** (two asterisks) for P < 0.01, *** (three asterisks) for P < 0.001, and **** (four asterisks) for P < 0.0001.

### eIF4G2 restrains a stemness-associated translational program with clinical relevance in PDA

To define the broader scope of eIF4G2-dependent translation, we profiled the whole-cell proteome of sg*Rosa* and sg*Eif4g2* PDA cells. Of the 79 transcripts whose translation was impaired in sg*Eif4g2* PDA cells, 31 (~40%) were detected by proteomic analysis and exhibited concordant decreases in protein abundance (Supplementary Table 3). Notably, 12 of these 31 (~40%) belong to a stemness-associated signature recently linked to eIF4G2 activity in intestinal stem cells (33) (Supplementary Table 3). Although not canonical EMT drivers, their combined loss would be expected to compromise epithelial integrity, deregulate polarity, and promote transcriptional reprogramming toward stem-like states. Among these, CREBBP is notable because it is frequently lost in human cancers (34–37). Indeed, we demonstrate that the heavy polysome association of the *Crebbp* mRNA (Supplementary Fig. 3J) and its protein expression (Fig. 3F) are strongly dependent on eIF4G2 expression. Consistent with the transcriptional functions of CREBBP, loss of eIF4G2 induces broad transcriptomic changes (Supplementary Fig. 3K, Supplementary Table 4) that are enriched for migration, wound healing, and neuronal features (Fig. 3G). In line with this, we observed robust upregulation of KRT14, a basal epithelial marker associated with poorly differentiated and stem-like PDA (7), in sg*Eif4g2* tumor lysates (Fig. 3H). Importantly, KRT14 induction was absent in sg*Pten* tumors (Supplementary Fig. 3L), supporting the notion that the stemness shift upon *Eif4g2* loss extends beyond PTEN suppression, reflecting a broader cell-state reprogramming program.

To contextualize these findings for human PDA, we projected the sg*Eif4g2* expression profile onto established human PDA transcriptional subtypes (6,38-40) and found strong enrichment for the “Basal” subtype (Fig. 4A), as well as EMT-related programs (Fig. 4B). In accordance with this observation, laser-capture microdissection of human PDA sections followed by RNA sequencing (19) revealed that *EIF4G2* mRNA is strongly depleted in poorly differentiated lesions (Supplementary Fig. 4A), providing a direct human correlate to our experimental observations. To further ascertain clinical relevance, we used VIPER (Virtual Inference of Protein activity by Enriched Regulon analysis) (41) with a curated *EIF4G2* regulon derived from our Riboseq and proteomic datasets to estimate eIF4G2 activity in human PDA transcriptomes. Inferred eIF4G2 activity was significantly reduced in metastases compared with primary tumors (Fig. 4C), and low activity correlated with poorer overall survival (Fig. 4D). Together, these results support a model in which eIF4G2 acts as a translational checkpoint linking selective mRNA control to PDA progression. By buffering against stemness-associated and EMT programs through regulators such as PTEN and CREBBP, eIF4G2 constrains metastatic plasticity, and loss of its activity marks a clinically aggressive subset of PDA tumors.

**Figure 4.**
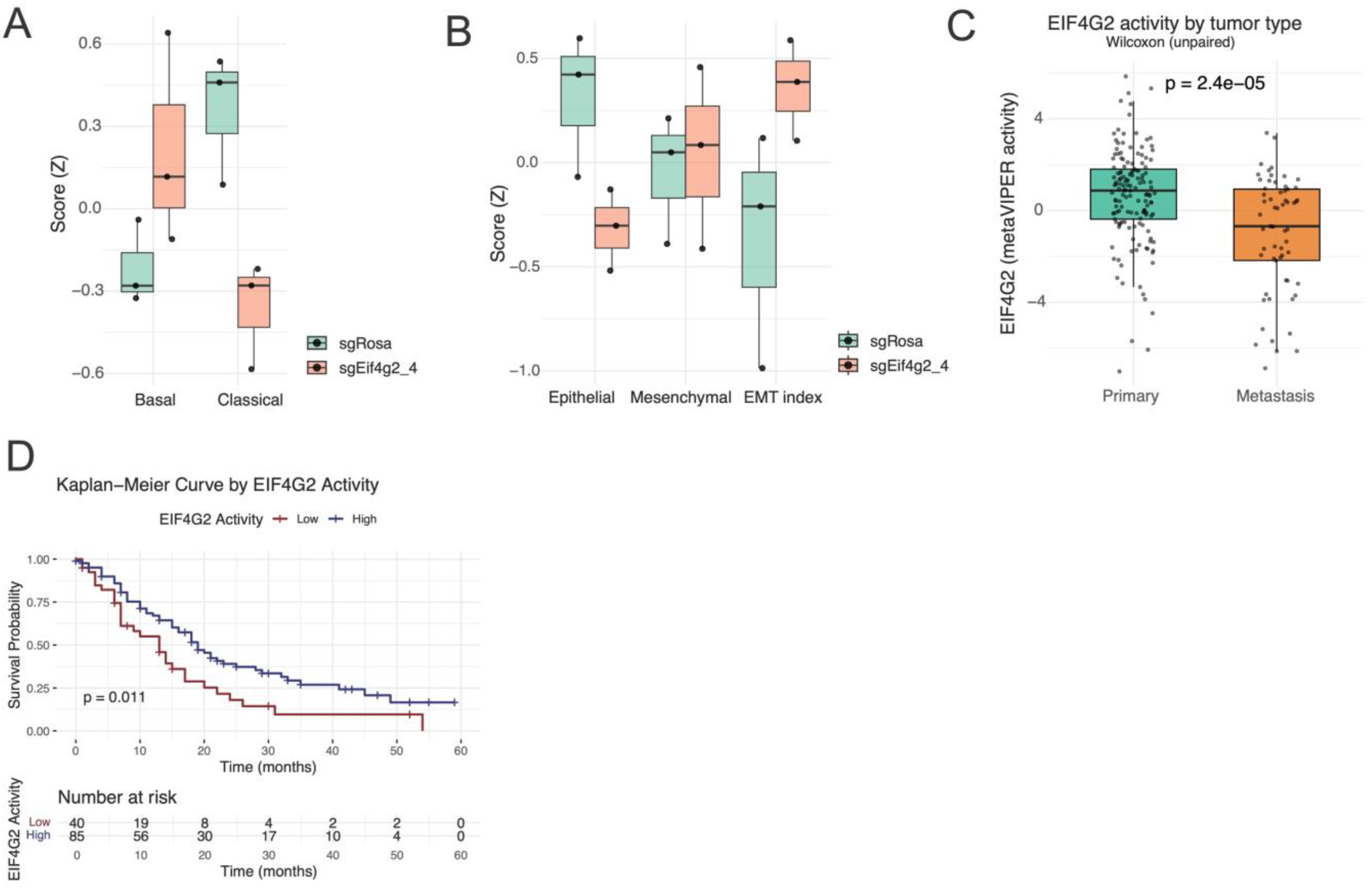
Loss of eIF4G2 promotes basal-like and metastasis-associated programs in human PDA. (A) Boxplots showing Z-score–based expression signature scores for Basal and Classical PDAC programs in control (sg*Rosa*) and EIF4G2 knockout (sg*Eif4g2*) cells. (B) Boxplots of Epithelial, Mesenchymal, and EMT index scores comparing control and eIF4G2 knockout cells. (C) EIF4G2 protein activity analysis in primary versus metastatic tumors from the UNC PDAC cohort (40). Differences between groups were evaluated using an unpaired, two-sided Wilcoxon rank-sum test; *p*-value indicated. (D) Kaplan-Meier analysis of overall survival of patients stratified by EIF4G2 activity (High > 0, Low ≤ 0) in the UNC PDAC cohort (40). Survival probability was estimated nonparametrically and compared with a two-sided log-rank test; the number at risk is shown at the bottom.

Collectively, our findings reveal that not all translational outputs drive oncogenesis; in PDA, eIF4G2-dependent translation actively restrains tumor progression. By sustaining translation of regulators such as PTEN and CREBBP, eIF4G2 maintains epithelial identity, and its loss drives dedifferentiation and metastatic spread. Reduced eIF4G2 activity in human metastases, together with its correlation with poor survival, highlights its potential as a prognostic biomarker. Its alignment with “Basal-like” features, including KRT14 induction, further supports integrating eIF4G2 activity scores into PDA subtype classifiers. Notably, because Basal-like PDA cells are more sensitive to KRAS inhibition (42), eIF4G2 activity scores may augment existing subtype classifiers and help stratify sensitivity to emerging KRAS-targeted therapies.

## MATERIALS AND METHODS

### Orthotopic allograft model of pancreatic cancer

Orthotopic engraftment of mouse pancreatic cancer cells was conducted as described (43). In brief, mice were anesthetized using isoflurane and subcutaneously administered 5 mg/kg Carprofen. 100,000 cells were transplanted to the parenchyma of the pancreas. The abdominal wall was sutured with absorbable Vicryl sutures (Ethicon Cat# J392H), and the skin was closed with wound clips (CellPoint Scientific Inc. Cat# 203-1000). C57Bl/6J mice (RRID:IMSR_JAX:000664) were purchased from the Jackson Laboratory for syngeneic orthotopic transplant experiments. All animal experiments were conducted in accordance with procedures approved by the IACUC at Columbia University (AC-AABK554).

### Cell culture conditions for monolayer cultures

Monolayer KPC primary pancreatic cancer cells were generated from tumors derived from *Trp53*^*+/LSL-R172H*^, *Kras*^*+/LSL-G12D*^, *Pdx1-Cre* (KPC) mice (44). HEK293T cell line was from ATCC (Cat# CRL-3216, RRID:CVCL_0063). All cells were cultured in DMEM (Gibco Cat#11995073) supplemented with 10% fetal bovine serum (FBS) (Corning Cat# 35-010-cv) and 1% PS unless stated otherwise. All cells were cultured at 37 °C with 5% CO_2_.

### CRISPR/Cas9-mediated gene deletion lentiviral production and transduction

KPC cells were transduced with lentivirus expressing Cas9 and sgRNAs (lentiCRISPRV2, Addgene # 52961). Single sgRNAs were cloned by annealing two DNA oligos and T4 DNA ligation into a BsmBI-digested lentiCRISPRV2. LentiCRISPRV2 lentiviruses were produced in HEK293T cells co-expressing the packaging vectors (pPAX2, Addgene # 12260 and VSV-G, Addgene # 12259), concentrated with LentiX concentrator (Clontech Cat# 631232), and resuspended with DMEM supplemented with 10% FBS and 1% PS at 10X concentration. Ecotropic pBabe-puro (Addgene Plasmid #1764) retroviruses were produced in Phoenix-ECO cells (ATCC Cat# CRL-3214), concentrated with RetroX Concentrator (Clontech Cat# 631456), and resuspended in DMEM supplemented with 10% FBS and 1% PS at 10X concentration. 100,000 cells were plated and infected with 5X concentrated viruses and spinoculated at 600 × *g* for 45 min at room temperature. One day after infection, cells were treated with 3 µg/mL puromycin (Sigma-Aldrich Cat # P9620) for selection.

sgRNA sequences used:

sg*Eif4g2*-2 sense, 5′-CACCGGTTCAGATAGTCAGTCACAA-3′

sg*Eif4g2*-2 antisense, 5′-AAACTTGTGACTGACTATCTGAACC-3′

sg*Eif4g2*-4 sense, 5′-CACCGATTAGACCATGAACGAGCCA-3′

sg*Eif4g2*-4 antisense, 5′-AAACTGGCTCGTTCATGGTCTAATC-3′

sg*Pten*-1 sense, 5′-CACCGACTATTCCAATGTTCAGTGG-3′

sg*Pten*-1 antisense, 5′-AAACCCACTGAACATTGGAATAGTC-3′

sg*Pten*-2 sense, 5′-CACCGGGTTTGATAAGTTCTAGCTG-3′

sg*Pten*-2 antisense, 5′-AAACCAGCTAGAACTTATCAAACCC-3′

sg*Rosa* sense 5′-CACCGAAGATGGGCGGGAGTCTTC-3′

sg*Rosa* antisense, 5′-AAACGAAGACTCCCGCCCATCTTC-3′

### CRISPR screen and analysis

#### CRISPR Library and Lentivirus Production

A genome-wide knockout screen was performed using the Brie mouse whole-genome CRISPR knockout library, which contains 78,637 gRNAs targeting 19,674 genes. For lentivirus production, 1.8 × 10^7^ HEK293T cells were seeded in a 175 cm^2^ tissue culture flask and the transfection was performed using a DNA mixture of VSV-G (5 µg), psPAX2 (50 µg), and 40 µg of the transfer vector. Flasks were transferred to a 37 °C incubator for 6–8 hours; after this, the media was aspirated and replaced with BSA-supplemented media. Virus was harvested 36 hours after this media change, pooled, and filtered.

#### Genome-wide CRISPR-Cas9 Knockout Screen

100 million ccmT2 cells, a monoclonal murine KPC PDA cell line, were transduced with the Brie lentiviral library at a multiplicity of infection (MOI) < 0.6. Three days post-transduction, the cells underwent selection with puromycin at 2 µg/mL for 3 days. After selection, a baseline cell sample was collected (Day 0). The remaining puromycin-selected cells were passaged in vitro for 17 days to deplete cells with knockouts of common essential genes. A sample of these “engraftment-ready” cells was collected (Day 17). For the in vivo screen, 1 x 10^6^ engraftment-ready cells were injected subcutaneously into the bilateral hind limbs of 10-week-old male B6J mice. The experiment was conducted with five cages of mice, each containing five male mice. Tumors were allowed to grow for 10 days, at which point they were harvested when they reached an average volume of approximately 1.5 cm^3^. Genomic DNA was isolated from the Day 0, Day 17, and endpoint tumor samples using the QIAamp DNA Mini Kit (Qiagen) according to the manufacturer’s instructions. The gDNA from 5 mice in the same cage were combined as one sample. The gRNA cassettes were amplified from the genomic DNA via a two-step PCR protocol. For the first step PCR, 500 µg of gDNA were used as input for one PCR reaction and in total 400 mg of gDNA were used for one cage of mice. The resulting amplicons were purified and subjected to deep sequencing on an Illumina NextSeq 500 platform to quantify gRNA abundance.

#### Data Processing and Analysis

Sequencing reads were processed and analyzed using the MAGeCK (v0.5.9) computational pipeline. Quality control analyses confirmed sufficient gRNA representation, high read-mapping rates (average > 85%), and uniform read distribution across all samples. Gene essentiality scores (beta scores) for the in vitro portion of the screen (Day 17 vs. Day 0) were calculated using MAGeCK-MLE and correlated with pan-cancer essentiality scores from the DepMap database (2025Q2). Gene-level depletion or enrichment in the in vivo screen (endpoint tumors vs. Day 17) was analyzed using MAGeCK-RRA. The results were compared with a previously published in vivo screen (22) to confirm technical robustness. All correlation analyses were performed using the Pearson correlation coefficient.

### Cell proliferation assays

#### CellTitre-Glo

Cell proliferation assay was performed by seeding 3,000 pancreatic cancer cells per well in opaque 96-well plates (Corning Cat# 3917). Cells were seeded in Gibco Fluorobrite DMEM (ThermoFisher Cat# A1896701) supplemented with 10% FBS, 1% P/S and 1% GlutaMAX (ThermoFisher Cat# 35050061). Cell viability was measured using a luminescent ATP-based assay (CellTiter-Glo, Promega Cat# G7573) with a plate reader (SpectraMax i3x, Molecular Devices). Data were analysed with GraphPad Prism.

#### EdU incorporation

EdU incorporation was assessed using the Click-iT™ EdU Alexa Fluor™ 488 Flow Cytometry Assay Kit (Thermo Fisher Scientific). Cells were treated with 10 µM EdU for 2 hours at 37°C, then trypsinized and fixed in 200 µL of 1% PFA for 10 minutes at room temperature with gentle rocking. After two washes with flow buffer (PBS + 1% BSA), cells were permeabilized in 200 µL of 1× BD Perm Buffer for 15 minutes at room temperature. Following two additional washes, cells were incubated in 100 µL of freshly prepared Click-iT™ reaction cocktail (430 µL reaction buffer, 20 µL CuSO_4_, 1.2 µL Alexa Fluor 488 azide, and 50 µL buffer additive) for 30 minutes at room temperature protected from light. Cells were then washed twice and analyzed by flow cytometry (Attune NxT, Thermo Fisher Scientific).

### Western blot analysis

Standard techniques were employed for immunoblotting of organoids. Protein lysates were prepared using 0.1% SDS lysis buffer in 50 mM Tris pH 8, 0.5% Deoxycholate, 150 mM NaCl, 2 mM EDTA, 1% NP40, with 1 tablet of PhosSTOP (Roche Cat# 4906837001) and 1 tablet of cOmplete™, Mini, EDTA-free Protease Inhibitor Cocktail (Roche Cat# 11836170001) per 10 ml buffer, and separated on 4-12% Bis-Tris NuPAGE gels (Invitrogen Cat# NP0335BOX) or 12% Bis-Tris SurePAGE Gel (Genscript Cat#M00669; M00667) or 3-8% Tris-Acetate NuPAGE gels (Invitrogen Cat# EA0378BOX), transferred onto a PVDF membrane (Millipore Cat# IPVH00010) and incubated with the indicated antibodies for immunoblotting. The following primary antibodies were used at 1:1000 dilution: eIF4G2 (D88B6) [Cell Signaling #5169, AB_10622189], eIF4A (C32B4)[Cell Signaling# 2013, AB_2097363], 4EBP1[Cell Signaling# 9452, AB_331692], pS6(D68F8)[Cell Signaling # 5364, AB_10694233], S6 (54D2) [Cell Signaling # 2317, AB_2238583], eIF4E(C46H6) [Cell Signaling # 2067, AB_2097675], PTEN (138G6) [Cell Signaling # 9559, AB_390810], CREBBP(D6C5)[Cell Signaling # 7389, AB_2616020].

### Proteomic Analysis

#### Global lysate proteome sample preparation

For each sample, 50 µg of protein were reduced with 5 mM dithiothreitol for 30 minutes at 25°C and 1,000 rpm, then alkylated in the dark with 10 mM iodoacetamide for 45 minutes at 25 °C and 1,000 rpm. Samples were then diluted with 50 mM Tris for a final urea concentration of < 2M. EDTA was added for a final concentration of 10 mM; lastly, SDS was added for a final concentration of 1%. Magnetic SP3 beads were made by combining equal volumes of carboxylate-modified hydrophilic (Cytiva: 45152105050250) and hydrophobic beads (Cytiva: 65152105050250). 500 µg of SP3 beads were added to each sample (input, flowthrough and IP). 100% ethanol was added at a 1:1 volumetric ratio with the sample to precipitate the protein material onto the beads. The samples were then incubated for 15 minutes at room temperature. Following incubation, the beads were washed thrice with 1 mL of 80% ethanol and reconstituted in 100 µL of freshly prepared ammonium bicarbonate with 0.5 µg of trypsin. The samples were incubated overnight at 37 °C degrees and 700 rpm to digest the proteins off of the SP3 beads. Tryptic peptides were then dried in a vacuum concentrator and resuspended in 3% acetonitrile/0.2% formic acid for a final peptide concentration of 0.25 µg/µL.

#### LC-MS/MS analysis on a Q-Exactive HF

Approximately 1 μg of total peptides were analyzed on a Waters M-Class UPLC using a 15 cm × 75 µm IonOpticks C18 1.7 µm column coupled to a benchtop Thermo Fisher Scientific Orbitrap Q Exactive HF mass spectrometer. Peptides were separated at a flow rate of 400 nL/min with a 150-minute gradient, including sample loading and column equilibration times. Data were acquired in data-independent mode using Xcalibur software. MS1 spectra were measured with a resolution of 120,000, an AGC target of 3e6 and a scan range from 350 to 1600 m/z. 35 isolation windows of 36.0 m/z were measured with a resolution of 30,000, an AGC target of 3e6, normalized collision energies of 22.5, 25, and 27.5, and a fixed first mass of 200 m/z.

Raw data were searched against the mus musculus proteome (UP000000589) with Spectronaut (version 19.4) and the DirectDIA workflow. The Biognosys Global Standard (BGS) factory settings were used, meaning cross-run normalization was implemented under the “Automatic” setting (RT-dependent local regression) and no imputation strategy was used. To measure relative protein abundance, iBAQ values were used as reported by Spectronaut. No further normalization was applied.

### Global translatome analysis

For each starting sample, cells were lysed and 1 µg of total RNA was isolated for RNA-Seq while 9 µg of total RNA was digested with RNase I to isolate ribosome protected RNA footprints (RPFs). Both sample types were taken through rRNA depletion and RNA-seq samples were heat fragmented. Both sample types then underwent adapter ligation. Ligated RNA was reverse transcribed, libraries were generated via PCR, and then sequenced on a NextSeq 2000. After sequencing, samples were processed with Eclipsebio’s proprietary analysis pipeline (v1). Unique molecular identifiers (UMIs) were pruned from read sequences using UMI-tools (v1.1.1). Next 3’ adapters were trimmed from reads using Cutadapt (v3.2). Reads were then mapped to a custom, curated database of repetitive elements and rRNA sequences. All non-repeat mapped reads were mapped to the genome hg38 using STAR (v2.7.7a), during the alignment multimapping reads were not retained. The exclusion of multimapping reads reduces false positive differential and pausing calls. The removal of multimapping reads may lead to underestimates of counts for genes with high rates of pseudogenization or duplication events. PCR duplicates were removed using UMI-tools (v1.1.1). Gene coverage was calculated using Eclipsebio’s proprietary feature counting algorithm and ribosome occupancy was calculated by dividing coverage in the ribosome protected footprint (RPF) library by the RNA-Seq library. Differentially expressed and occupied genes were identified using DESeq2. For transcriptome analyses, genomic aligned reads were aligned to the transcriptome with mudskipper (v0.1.0). Transcriptome aligned reads were processed with riboWaltz and custom analysis programs developed by Eclipsebio to determine codon usage, stall sites, 5’ UTR loading, and 3’ read through. ORF detection was performed on genome alignments using ribotricer (v1.3.3).

### *O*-propargyl-puromycin labeling

Click-iT assays were performed using 1 × 10^6^ cells per assay according to the manufacturer’s protocol. In brief, *O*-propargyl-puromycin (OP-Puro; 20 μM) (Life Technologies Cat# C10456,) was added to the cells and incubated for 30 minutes. Cells were washed in ice-cold PBS and then fixed and permeabilized. The amount of Alexa-Fluor-488 conjugated OP-Puro was quantified using flow cytometry using a BD LSR Fortessa Cytometer (BD Biosciences). Data analysis was performed using FlowJo v9.7.6 (FlowJo, Ashland, RRID:SCR_008520).

### Polysome fractionation and qRT-PCR

100 μg/ml of cycloheximide was supplemented into the media. The cells were then harvested on ice in PBS containing 100 μg/ml cycloheximide. Cells were pelleted and lysed in 10 mM Tris-HCl (pH 8), 140 mM NaCl, 1.5 mM MgCl_2_, 0.25% NP-40, 0.1% Triton X-100, 50 µM DTT, 150 μg/mL cycloheximide, and 640 U/mL RNaseIN (Sigma Cat# 3335399001) for 15 minutes. Lysates were cleared and then loaded onto a 10-50% sucrose gradient made using a Biocomp Gradient Master 108 and centrifuged for 2 hours and 15 minutes at 35,000 rpm in a SW41 rotor using a Sorvall Discovery 90SE. The gradients were fractionated on a Teledyne ISCO Foxy R1 apparatus while monitoring the OD_254_. The following SybrGreen primers were used:

*Pten*: Forward 5’-GAAAGGGACGGACTGGTGTA, Reverse 5’-TAGGGCCTCTTGTGCCTTTA

*Crebbp*: Forward 5’-CACCATCTGTGGCTACTCCTCA, Reverse 5’-GGTTTCAGCACTGGTCACAGAG

### L-Azidohomoalanine (AHA)-based detection of nascent protein synthesis by immunoblot

Cells were incubated in methionine-free medium at 37 °C for 30 minutes to deplete intracellular methionine. Nascent protein synthesis was then pulse-labeled by adding AHA or methionine (negative control) to the methionine-free medium and incubating for at least 10 minutes at 37 °C. Cycloheximide was used in parallel as a translational inhibition control. Following treatment, cells were washed with ice-cold PBS and lysed in RIPA buffer lacking EDTA (containing Tris pH 8.0, NaCl, NP-40, sodium deoxycholate, and SDS). Protein concentration was measured using a BCA assay. Equal amounts of total protein were prepared in Click-iT™ reaction buffer and incubated with a freshly prepared cocktail containing CuSO_4_, biotin-alkyne, and Click-iT™ buffer additive. Samples were incubated at room temperature to allow copper-catalyzed azide–alkyne cycloaddition (CuAAC). Proteins were separated by SDS-PAGE and transferred to nitrocellulose membranes. Biotin-labeled proteins were detected by probing with streptavidin–HRP (Cell Signaling Cat #3999) at a 1:1000 dilution and visualized by chemiluminescence.

### Histological annotation and quantification of tumors

Whole slide imaging of H&E (hematoxylin and eosin)-stained tumor sections was performed using the Aperio AT2 slide scanner (Leica), and the images were analyzed using QuPath software. For each sample, regions of well-, moderately-, and poorly differentiated morphology were annotated based on standard histopathological criteria by an experienced pathologist. The percentage of each category was quantified relative to the total annotated tumor area per slide, with values summing to 100% per sample. All annotations were performed blinded to sample identity. To enable subtype-relevant comparisons, a secondary classification was applied using a glandular architecture-based system previously shown to correlate with transcriptional subtypes of pancreatic ductal adenocarcinoma (PDAC) (45). In this schema, gland-forming regions are enriched in Classical subtype tumors, whereas non-gland-forming regions are characteristic of Basal-like PDAC. All well-differentiated areas were gland-forming, and all poorly differentiated areas were non-gland-forming. Moderately differentiated areas were further subclassified into gland-forming and non-gland-forming categories based on the presence or absence of defined glandular architecture. Quantitative measurements of glandular and non–glandular composition were generated accordingly, offering a subtype-informed stratification of tumor differentiation status.

### eIF4G2 activity analysis on metastasis and survival outcomes

Expression data and clinicopathological metadata, including sample type (“Primary” vs. “Metastasis”) and epithelial subtype (basal, classical, precursor), were obtained from the UNC pancreatic ductal adenocarcinoma cohort (40). Protein activity was inferred using VIPER (41) applied to expression signatures. EIF4G2 activity values were extracted from the resulting activity matrix, and the activity difference between primary and metastatic tumors was evaluated with an unpaired, two-sided Wilcoxon rank-sum test. Since this was a single, pre-specified comparison, no multiple-testing correction was applied. Analyses were performed in R (version ≥4.3), and boxplots were generated using ggplot2. For survival analysis, samples were divided into two groups (EIF4G2 High: activity > 0; EIF4G2 Low: activity ≤ 0), a threshold that corresponds to the sign change of the standardized VIPER score. Overall survival was defined as the time in months from diagnosis (or the cohort-defined index) to death. Kaplan–Meier survival curves were estimated nonparametrically, and differences between groups were compared using a two-sided log-rank (Mantel–Cox) test with α = 0.05. Analyses were performed in R (version ≥4.3) using the survival package to create Survival objects, fit Kaplan–Meier estimators, and perform log-rank tests, and with survival for visualization with number-at-risk tables. Greenwood standard errors were used by default, and unadjusted log-rank P values are reported.

### eIF4G2 z-score analysis on basal/classical/Epithelial/EMT signatures

Bulk RNA-seq counts from control (sgROSA) and EIF4G2 knockout (sgEIF4G2) cells were analyzed with DESeq2 (46). For signature scoring, variance-stabilized expression values were calculated and converted to per-gene Z-scores by centering and scaling across all samples. Gene sets included (i) Moffitt epithelial programs defining Classical and Basal-like PDAC (40), and (ii) curated epithelial and mesenchymal marker panels for EMT (47). Signature scores were obtained for each sample as the mean Z-score of genes from the respective set present in the expression matrix. An EMT index was calculated as (Mesenchymal score – Epithelial score). Marks used for each analysis are listed below:

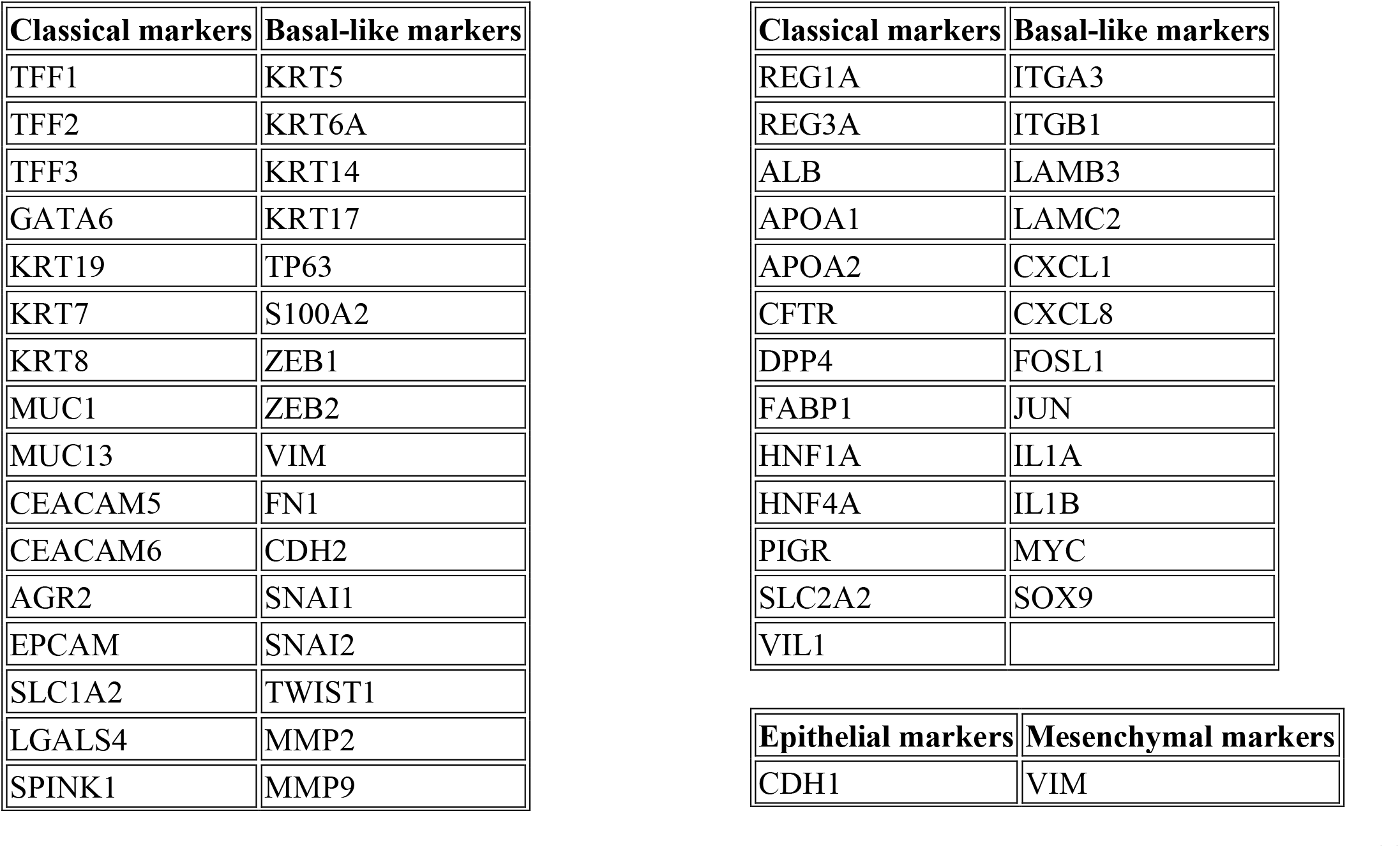
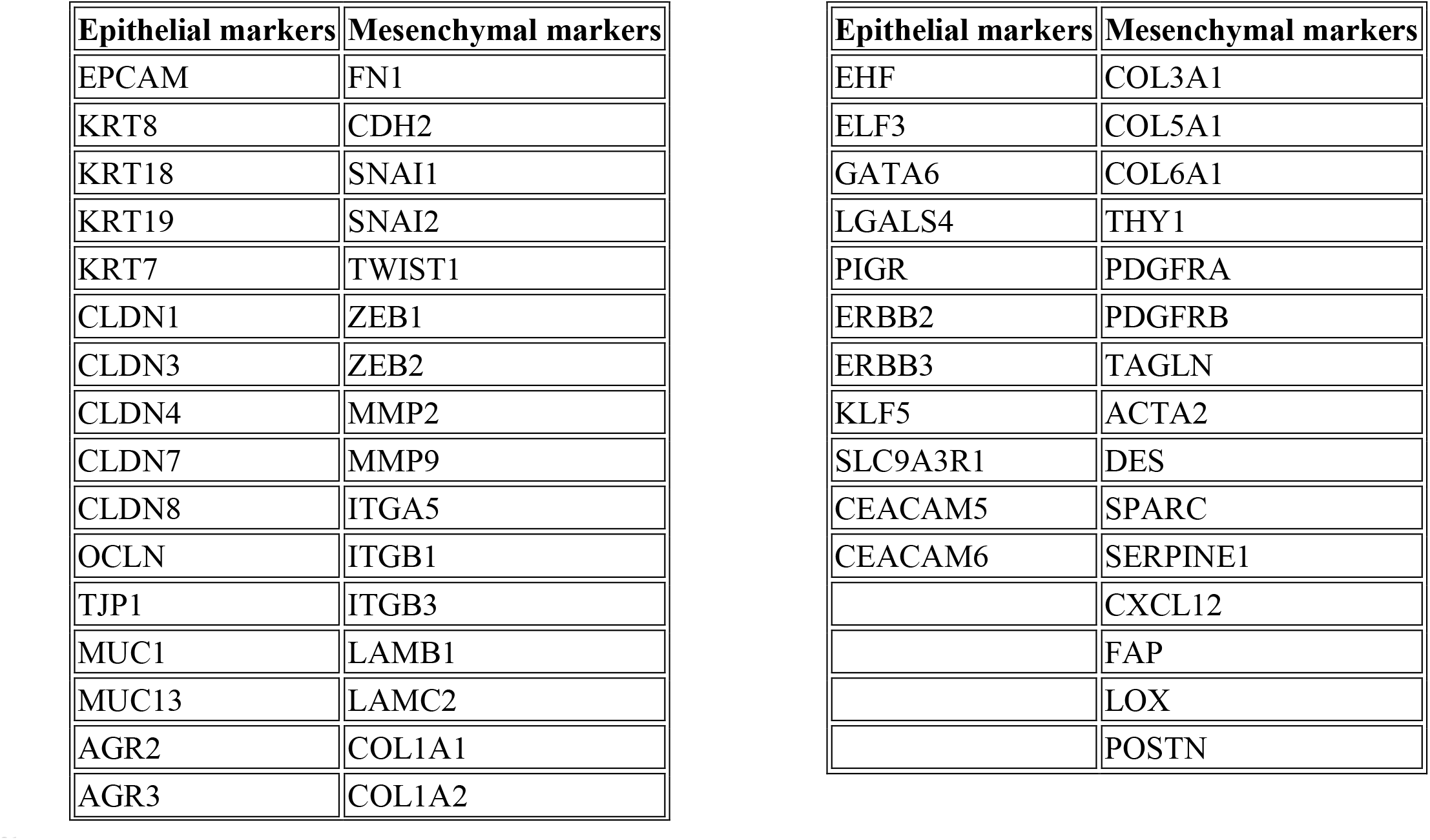

| Classical markers | Basal-like markers |
| --- | --- |
| TFF1 | KRT5 |
| TFF2 | KRT6A |
| TFF3 | KRT14 |
| GATA6 | KRT17 |
| KRT19 | TP63 |
| KRT7 | S100A2 |
| KRT8 | ZEB1 |
| MUC1 | ZEB2 |
| MUC13 | VIM |
| CEACAM5 | FN1 |
| CEACAM6 | CDH2 |
| AGR2 | SNAI1 |
| EPCAM | SNAI2 |
| SLC1A2 | TWIST1 |
| LGALS4 | MMP2 |
| SPINK1 | MMP9 |
| REG1A | ITGA3 |
| REG3A | ITGB1 |
| ALB | LAMB3 |
| APOA1 | LAMC2 |
| APOA2 | CXCL1 |
| CFTR | CXCL8 |
| DPP4 | FOSL1 |
| FABP1 | JUN |
| HNF1A | IL1A |
| HNF4A | IL1B |
| PIGR | MYC |
| SLC2A2 | SOX9 |
| VIL1 |  |

| Epithelial markers | Mesenchymal markers |
| --- | --- |
| CDH1 | VIM |
| EPCAM | FN1 |
| KRT8 | CDH2 |
| KRT18 | SNAI1 |
| KRT19 | SNAI2 |
| KRT7 | TWIST1 |
| CLDN1 | ZEB1 |
| CLDN3 | ZEB2 |
| CLDN4 | MMP2 |
| CLDN7 | MMP9 |
| CLDN8 | ITGA5 |
| OCLN | ITGB1 |
| TJP1 | ITGB3 |
| MUC1 | LAMB1 |
| MUC13 | LAMC2 |
| AGR2 | COL1A1 |
| AGR3 | COL1A2 |
| EHF | COL3A1 |
| ELF3 | COL5A1 |
| GATA6 | COL6A1 |
| LGALS4 | THY1 |
| PIGR | PDGFRA |
| ERBB2 | PDGFRB |
| ERBB3 | TAGLN |
| KLF5 | ACTA2 |
| SLC9A3R1 | DES |
| CEACAM5 | SPARC |
| CEACAM6 | SERPINE1 |
|  | CXCL12 |
|  | FAP |
|  | LOX |
|  | POSTN |

### Quantifications and Statistical Analysis

Biochemical experiments *in vitro* were repeated at least three times, and the repeat number was increased according to effect size or sample variation. We estimated the sample size considering the variation and mean of the samples. No statistical method was used to predetermine the sample size. No animals or samples were excluded from any analysis. Animals were randomly assigned groups for *in vivo* studies; no formal randomization method was applied when assigning animals for treatment. All western blotting experiments with quantification were performed a minimum of three times with independent biological samples and analyzed by ImageJ 1.52q. Investigators were blinded to group allocation during data analysis. Statistical analyses were performed using GraphPad Prism 8. All tests and *p* values are provided in the corresponding Figs. or Fig. legends.

## AUTHORS’ CONTRIBUTIONS

**J. Powers**: Conceptualization, data curation, methodology, writing—review and editing. **W. Lai:** Conceptualization, formal analysis, methodology, writing—original draft, writing—review and editing. **P. Ahmadi:** Data curation, writing—review and editing. **H. Kobayashi:** formal analysis, methodology, writing— original draft, writing—review and editing. **A. Curiel-Garcia:** formal analysis, methodology, writing— original draft, writing—review and editing. **E. Valenzuela**: data curation, formal analysis, methodology, writing—original draft, writing—review and editing. **M. Jovanovic**: formal analysis, methodology, writing— original draft, writing—review and editing. **A. Chavez**: Conceptualization, formal analysis, methodology, writing—original draft, writing—review and editing. **I.I.C. Chio**: Conceptualization, resources, data curation, formal analysis, supervision, funding acquisition, investigation, project administration, writing—original draft, writing–review and editing.

## ACKNOWLEDGEMENTS

We thank other members of the Chio lab, as well as Drs. Richard Baer, Laura Pasqualucci, and Luke Berchowitz for discussion. This work was performed with the support of the Herbert Irving Comprehensive Cancer Center (Columbia University Irving Medical Center) Proteomics, Flow Cytometry, Genomics and High Throughput Screening, Oncology Precision Therapeutics and Imaging Core, and Molecular Pathology (MPSR) Shared Resources, as well as the Mass Spectrometry Core Facility (Chemistry Department at Columbia University).

## SUPPLEMENTARY FIGURES

**Supplementary Figure 1.**
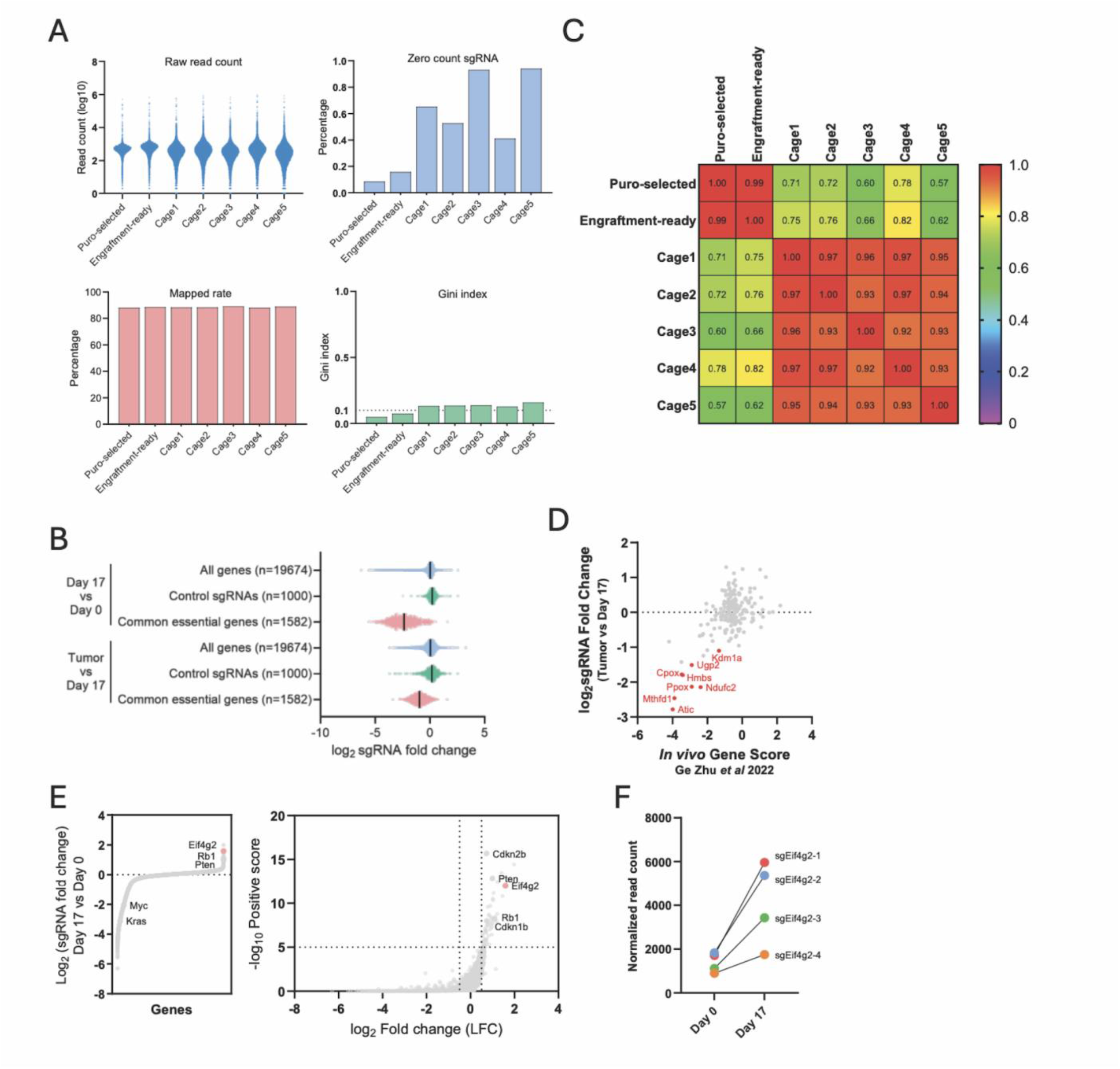
Quality assessment and validation of the genome-wide CRISPR screen. (A) Distribution of log-transformed sgRNA read counts. Other metrics for screen quality are also shown such as read mapping rates (average > 85%), sgRNA dropout, and uniform read distribution (average Gini index ≈ 0.1). (B) Distribution of sgRNA log2-fold change for the in vitro (Day 17 vs. Day 0) and *in vivo* (endpoint tumors vs. Day 17) portions of the screen. (C) Correlation matrix of sgRNA read counts across different screening conditions. (D) The correlation of the sgRNA fold change from our in vivo screen with the gene score generated from the PDA *in vivo* screen performed in the Birsoy lab (22). (E) The fold change of sgRNAs and volcano plot of the in vitro screen results (Day 17 vs. Day 0), the enrichment of *Eif4g2* is highlighted. (F) Normalized read counts for four independent sgRNAs targeting *Eif4g2*, between the Day 0 baseline and the Day 17 cell population.

**Supplementary Figure 2.**
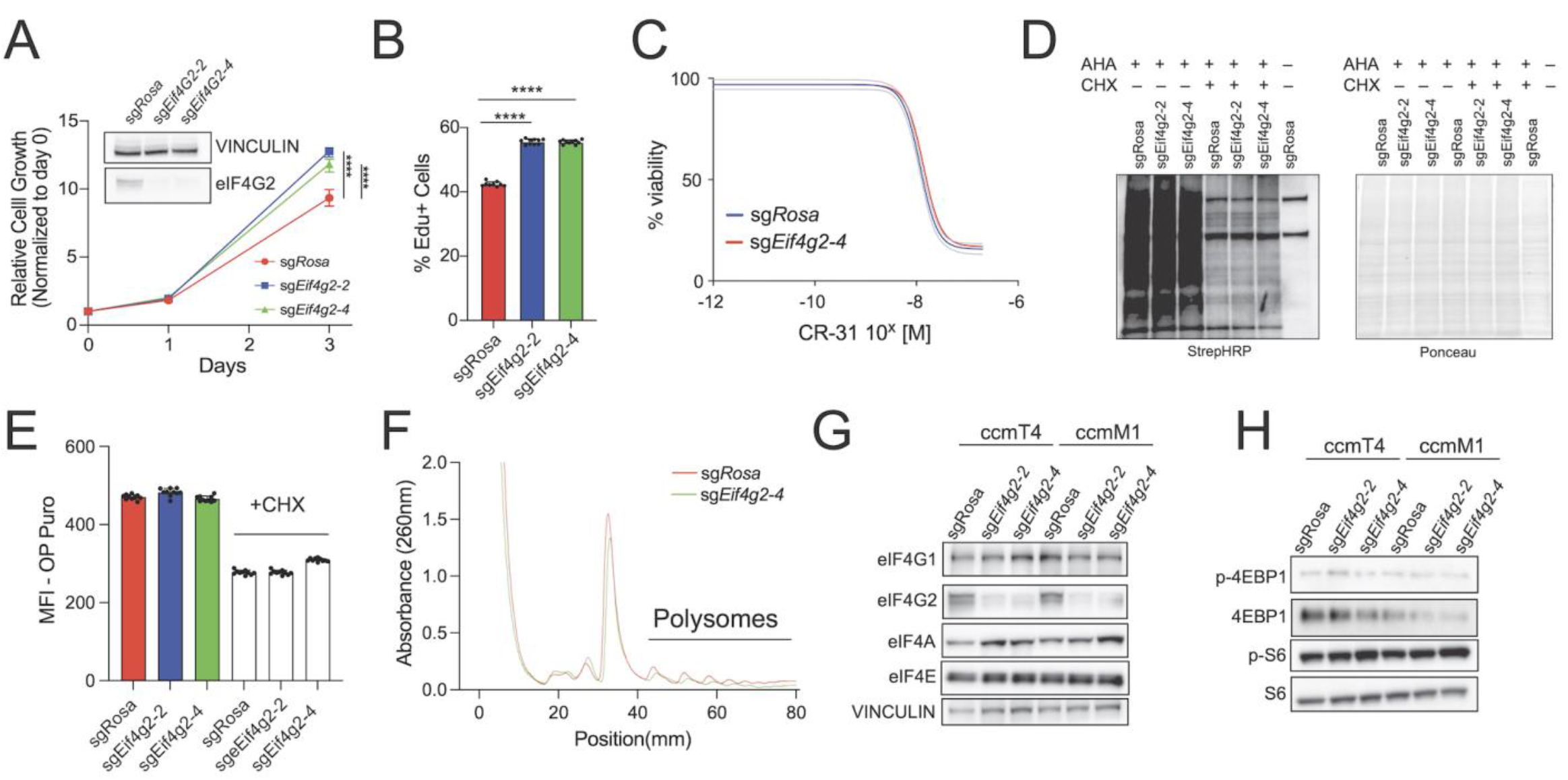
Loss of eIF4G2 does not alter global protein synthesis. (A, B) Proliferation of control (sg*Rosa*) and eIF4G2-deficient (sg*Eif4g2*) KPC cells measured by CellTiter-Glo (A) and EdU incorporation (B). (C) Viability of control and eIF4G2-deficient KPC cells upon increasing concentrations of eIF4A inhibitor (CR-1-31-B) for 72 hours. (D) L-Azidohomoalanine (AHA) incorporation in sg*Rosa* and sg*Eif4g2* KPC cells was analyzed by immunoblot, with cells treated with cycloheximide serving as negative controls. (E) OP-Puro incorporation was similarly analyzed by immunoblot, with cycloheximide-treated cells as controls. (F)Polysome profiles of control and eIF4G2-deficient KPC cells. Absorbance at 254 nm is plotted as a function of sedimentation. (G and H) Representative immunoblot analysis of the abundance of eIF4F components (I) and downstream substrates of mTOR activation (J). Error bars in this Fig. are means ± SDs. Student’s t-test was performed. ns (not significant) for P ≥ 0.05, * (one asterisk) for P < 0.05, ** (two asterisks) for P < 0.01, *** (three asterisks) for P < 0.001, and **** (four asterisks) for P < 0.0001.

**Supplementary Figure 3.**
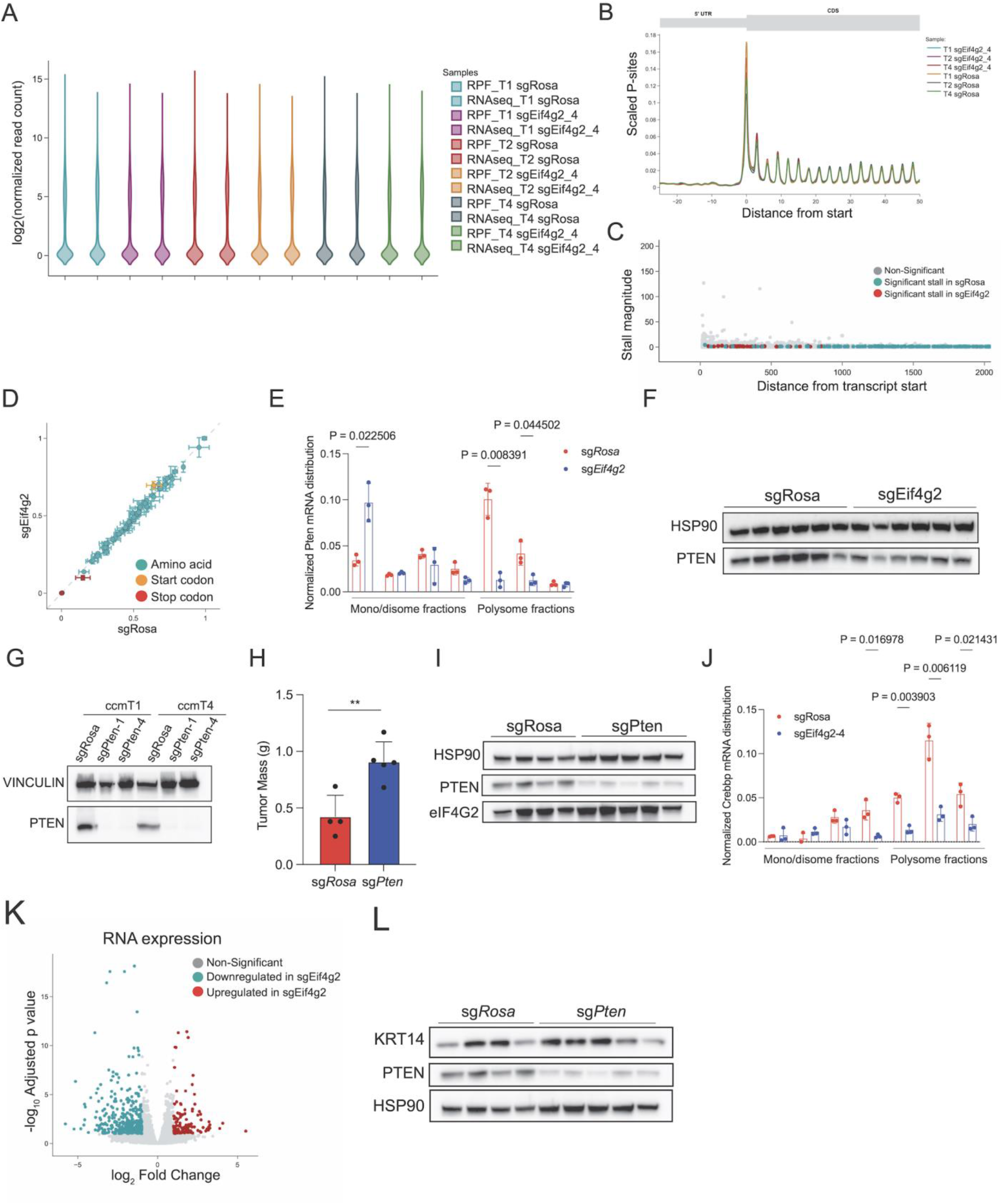
Ribosome profiling to characterize eIF4G2-dependent translatome. (A) Pairs of ribosome-protected footprint (RPF) and RNA-Seq libraries were prepared from each sample. The plot shows the distribution of RPM (reads per million) normalized gene counts in each of the libraries. (B) This plot shows the location of scaled read ends that have been corrected for their estimated P-site (the second binding site for tRNA in the ribosome) from the ribosome-protected footprint (RPF) libraries. This correction is to show where ribosomes are directly binding, as the sequenced read ends have an overhang past the ribosome complex. RPF libraries are expected to show a trinucleotide periodicity. (C) RPF reads will accumulate where a ribosome pauses or stalls during translation. This plot shows putative stall locations for each transcript in the transcriptome, where stalls were detected through the use of a Kolmogorov-Smirnov (KS) test on normalized cumulative P-site densities. This analysis is performed per transcript, so the listed distances refer to the distance in transcriptomic space, not genomic space. (D) This scatter plot shows the mean codon usage in each condition, where codon usage is defined as the scaled normalized frequency that P-sites were found to be enriched in each codon. Each dot is a different codon, colored by whether it is a start, stop, or an amino acid-coding codon. Off-diagonal codons can indicate differential codon usage between the two conditions. Codon usage was determined using riboWaltz. If a gene has multiple transcripts, only the longest transcript is included in this analysis. (E) qRT-PCR analysis of *Pten* mRNA distribution across polysome fractions. (F) immunoblot analysis of PTEN expression in control (sg*Rosa*) and eIF4G2-deficient (sg*Eif4g2*) bulk tumor lysates. (G) Immunoblot analysis of PTEN expression in sg*Rosa* and sg*Pten* KPC cells. (H) Mass of pancreatic tumors from in sg*Rosa* and sg*Pten* KPC cells 5 weeks after orthotopic implantation. (I) Immunoblot analysis of eIF4G2 expression in control (sg*Rosa*) and *Pten*-deficient (sg*Pten*) bulk tumor lysates. (J) qRT-PCR analysis of *Crebbp* mRNA distribution across polysome fractions. (K) Volcano plot of eIF4G2-dependent transcriptional changes. (L) KRT14 and PTEN expression in sg*Rosa* and sg*Pten* bulk tumor lysates. PTEN and HSP90 immunoblots are also used in panel (I). Error bars in this Fig. are means ± SDs. Student’s t-test was performed. ns (not significant) for P ≥ 0.05, * (one asterisk) for P < 0.05, ** (two asterisks) for P < 0.01, *** (three asterisks) for P < 0.001, and **** (four asterisks) for P < 0.0001.

**Supplementary Figure 4.**
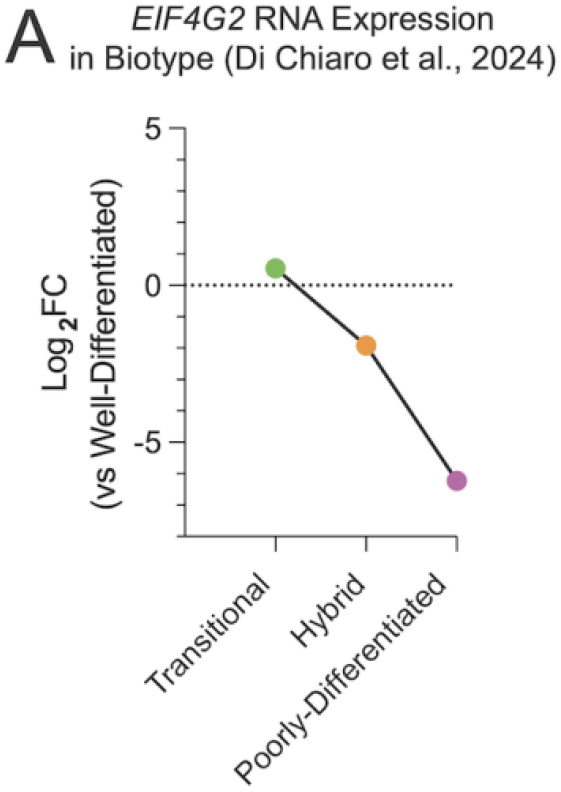
eIF4G2 expression in human PDA. (A) *EIF4G2* mRNA abundance based on a recently published morpho-biotype framework for PDA, relative to well-differentiated lesions.

**Supplementary Table 1**. Summary of metastatic lesions identified in each mouse transplanted with sg*Rosa* or sg*Eif4g2* PDA cells by organ site.

**Supplementary Table 2**. List of mRNAs that exhibit significantly different ribosomal occupancy, i.e., translation efficiency, in sg*Eif4g2* PDA cells compared to sg*Rosa* control PDA cells.

**Supplementary Table 3**. Translationally downregulated transcripts with matching protein decreases in sg*Eif4g2* vs sg*Rosa* PDA cells.

**Supplementary Table 4**. List of differentially expressed transcripts in sg*Eif4g2* PDA cells compared to sg*Rosa* control PDA cells.

